# Generalist large language models complement tailor-made predictors for tumor genomics interpretation

**DOI:** 10.64898/2026.05.21.726957

**Authors:** Jennifer Yu, Madison Darmofal, Michele Waters, January Choy, Thinh N. Tran, Chenlian Fu, Leah Morales, U Kaicheng, Ross L. Levine, Nikolaus Schultz, Michael F. Berger, Quaid Morris, Justin Jee

## Abstract

General-purpose large language models (LLMs) are trained on large corpora to acquire broad knowledge, but whether LLMs can replace, or augment, task-specific models is unclear. We evaluated LLMs on three real-world, clinically important tumor genomic interpretation tasks, in order of increasing difficulty: (i) distinguishing tumor from non-tumor mutations (n=34,415 variants), (ii) distinguishing driver from passenger mutations (n=13,469 variants), and (iii) inferring cancer type from tumor sequencing reports across multiple assays and institutions (n=102,791 samples). The best general-purpose LLMs performed as well as the benchmark tailor-made predictor for task (i). Ensembling tailor-made models with zero-shot LLMs improved their performance for tasks (i) and (ii). For task (iii), LLMs outperformed or supplemented tailor-made models on out-of-distribution data. Without fine-tuning, current LLMs already can be useful in clinical genomic interpretation by adding complementary expertise to tailor-made, state-of-the-art predictors.

## Main

Generalist large language models (LLMs) trained on large natural language corpora perform well at many prediction problems in science and medicine ^1,2^. It is tempting to postulate that LLMs are now poised to render obsolete the specialized medical models currently in use, the way they have in other fields, from playing chess to computer vision^3^.

Tumor genomic profiling is an integral part of cancer care^4^. However, interpreting clinical genomic reports is challenging for both patients and clinicians. Specially-trained bioinformatics algorithms are used in many aspects of interpretation, for example in distinguishing non-tumor from tumor mutations, distinguishing cancer driver mutations from passenger mutations, and inferring cancer type based on genomic alterations^5–7^. Performance aside, because of the relatively high barrier to implementation of specialized bioinformatics tools, it is plausible that patients and clinicians may use LLMs directly to help with tumor genomics interpretation. However, the extent to which LLMs can replace or augment boutique algorithms for tumor genomic interpretation has been mostly studied in limited cohorts or with oversimplified tasks^8,9^.

Leveraging large validation datasets, we evaluated the zero-shot performance of general-purpose proprietary, open weight, and domain-specialized medical LLMs on challenging, real-world tumor genomic tasks. We benchmarked their performance against tailor-made predictors. We further tested whether ensemble models incorporating zero-shot LLMs and tailor-made predictors could outperform either model in isolation.

### Tumor vs non-tumor mutations

To reduce costs and complexity, many clinical sequencing assays do not sequence matched blood and rely on computational methods to distinguish tumor-associated somatic variants from others. While it is often easy to identify germline-variants based on variant allele frequency (VAF), in lymphocyte-infiltrated tumors, variants associated with clonal hematopoiesis of indeterminate potential (CHIP) can reach detectable allele frequencies within the tumor sample and be incorrectly annotated as cancerous^10^. Using a cohort of patients^11^ with solid tumors and matched tumor and white blood cell (WBC) sequencing, which can be used to screen for CHIP^12^, we curated a set of CHIP and tumor somatic variants. We benchmarked zero-shot LLMs and MetaCH ^5^, a state-of-the-art supervised machine-learning framework trained on our curated cohort. To use the LLMs for zero-shot predictions, we prompted each with a structured mutation report containing gene, reference allele, tumor allele, chromosome and position, mutation type, VAF, and cancer type, thereby recapitulating the inputs to MetaCH. After excluding samples used for training MetaCH, N=34,415 variants from 16,383 samples (12,202 patients), were included in our analysis.

For somatic-vs-CHIP classification, Qwen3-30B achieved the highest overall accuracy among LLMs (89.9%) with per-cancer-type Q1–Q3: 86.6%–92.6% (Fig. 1a, Supplementary Fig. 1a, Supplementary Table 1–2). MetaCH’s overall accuracy was 89.7% (per-cancer-type Q1–Q3: 84.0%–91.9%). LLMs and MetaCH performed best at identifying CHIP variants in canonical CHIP genes (e.g., *DNMT3A, TET2, ASXL1*, Supplementary Fig. 2a). Ablating VAF significantly decreased performance across most models (Supplementary Fig. 3a), demonstrating the importance of allele frequency in distinguishing CHIP variants from tumor mutations.

**Figure 1.**
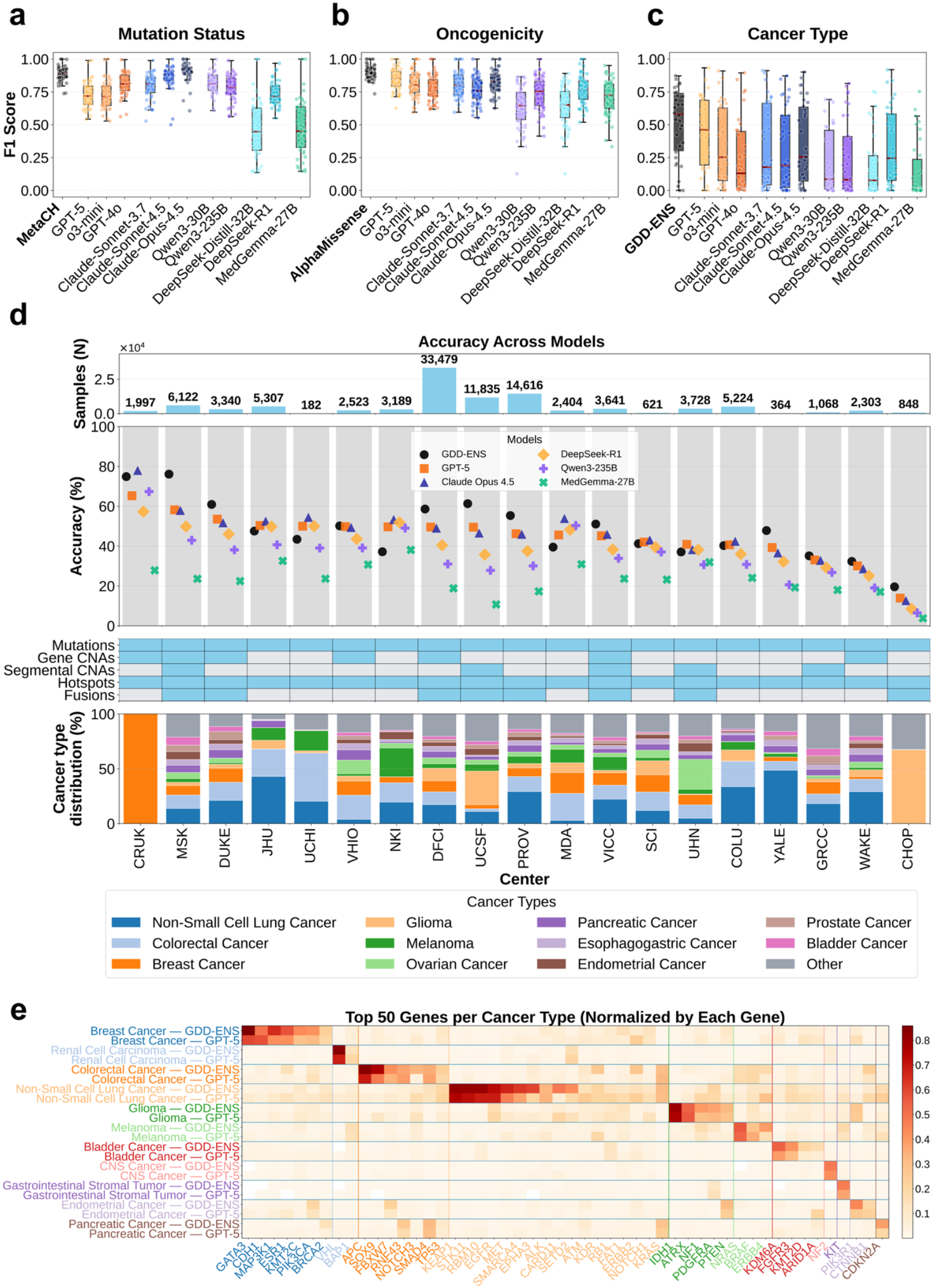
LLM performance across tumor genomic interpretation tasks. a, Mutation status classification (tumor somatic vs. CHIP). Box plots show F1 score distribution across cancer types for MetaCH and each LLM. b, Oncogenicity prediction compared with AlphaMissense. c, Cancer type prediction compared with GDD-ENS. Box plots show median (line), interquartile range (box), and 1.5x IQR (whiskers). Individual points represent cancer types. Red dashed line indicates the mean. d, Cancer type prediction accuracy across 19 GENIE centers. Top panel shows sample size per center. Middle panel shows accuracy for each model (GDD-ENS, GPT-5, Claude Opus 4.5, DeepSeek-R1, Qwen3-235B, MedGemma-27B). Bottom panels show data type distribution and cancer type composition per center. e, Normalized prediction probabilities for top 50 genes across cancer types, comparing GDD-ENS and GPT-5. Genes clustered by primary cancer association. Color intensity indicates prediction probability (0–1).

### Driver vs passenger mutations

Distinguishing driver from passenger mutations is important for prognostication and selecting therapies. We curated 13,469 variants (4,652 non-driver and 8,817 driver mutations) from 10,752 clinically sequenced tumor samples (10,489 patients), with reference labels derived from OncoKB^13^. Because variants of unknown significance (VUS) may include pathogenic alterations, for negative controls, as per prior work^14^, we randomly sampled benign germline missense variants, treating these as negative controls. As we previously reported that AlphaMissense^6^ – a deep learning model for proteome-wide missense effect prediction – was the best-in-class model for discriminating tumor driver and passenger mutations; we use it here as the benchmark. LLMs, particularly GPT-5 (overall accuracy 77.5%; per-cancer-type Q1–Q3: 79.6%– 90.5%), Claude-Opus-4.5 (75.1%; Q1–Q3: 76.6%–88.2%) and o3-mini (75.2%; Q1–Q3: 74.5%–88.0%) achieved accuracies approaching AlphaMissense (86.6%; per-cancer-type Q1–Q3: 86.2%–92.9%) (Fig. 1b, Supplementary Table 1–2) across cancer types and genes. (Supplementary Fig. 1b, Supplementary Fig. 2b).

It is possible that some VUSs may be drivers but lack experimental confirmation required for driver status in OncoKB and thus our positive control dataset. To further test the utility of AI-generated annotations among VUSs, we employed a recently-described approach to assess driver predictions using real-world survival data ^14^. Specifically, we accessed survival among non-small cell lung cancer patients stratified by mutations in *KEAP1*, in which known drivers are associated with worse prognosis. Patients with *KEAP1* VUSs annotated as oncogenic by LLMs had survival curves similar to those with *KEAP1* mutations, known to be oncogenic (HR 1.63, 95% CI 1.50-1.79); whereas patients with LLM-predicted benign VUSs had survival curves not significantly different than those without *KEAP1* mutations (HR 1.14, 95% CI 0.73 – 1.80). These results support the clinical validity of LLM-based oncogenic classifications of VUS (Supplementary Fig. 4).

### Cancer type inference from tumor genomics

Determining cancer type is a critical part of patient diagnosis and treatment planning. Genomic data can supplement inference from histopathologic slides, particularly in ambiguous cases or cases of cancer of unknown primary ^15^. This task is made challenging by the many possible genomic alterations and substantial overlap in certain alterations across cancer types. Nonetheless, machine learning algorithms, under supervised training with tens of thousands of examples, have shown promise at distinguishing cancer type using clinical genomic profiles alone ^7,15,16^. We evaluated whether zero-shot LLMs can infer cancer type from genomic alterations in a cohort of 102,791 samples from 97,074 patients across 19 institutions (the AACR Project GENIE dataset), spanning 34 cancer types. Molecular profiles could include somatic mutations, copy number alterations, arm-level chromosomal changes, or gene fusions. We benchmarked against GDD-ENS ^7^, a best-in-class supervised ensemble deep learning model trained for cancer type prediction from genomic features based on tumor genomic profiling data from Memorial Sloan Kettering (MSK).

GPT-5 was the best-performing LLM (overall accuracy 48.1%; per-cancer-type accuracy Q1– Q3: 10.7%–52.3%, Fig. 1c, Supplementary Table 1–2). GDD-ENS achieved an overall accuracy of 54.7% (per-cancer accuracy Q1–Q3: 17.9%–58.5%). Performance varied substantially across centers, which included different cancer-type distributions and sequencing assays (Fig. 1d). In centers with fewer genes included in their sequencing panel or missing modalities (e.g., gene-level or segmental copy number calls, fusions), LLMs performed better than GDD-ENS. These results suggest that GDD-ENS suffers when its trained feature set is incomplete, while LLMs can leverage incomplete data more flexibly. Performance was also notably low at the Children’s Hospital of Philadelphia (CHOP), whose cohort consisted almost entirely of gliomas (predominantly pediatric low-grade gliomas). This a population underrepresented in the GDD-ENS training set and intrinsically difficult to classify by all models.

To test whether LLMs identified features that were like those GDD-ENS identified as important, we prompted each LLM to report the three most influential genes driving its cancer-type prediction. LLM-identified influential features correlated well with SHAP feature importances from GDD-ENS (mean Pearson r = 0.902 across cancer types; mean r = 0.713 across genes), indicating reliance on clinically meaningful genomic signals. Both approaches recovered canonical gene–cancer associations (e.g., *GATA3*–breast^17^, *APC*–colorectal^18^, *KEAP1*/*STK11*/*EGFR*–non-small cell lung cancer^19^, Fig. 1e; Supplementary Fig. 3b–c).

### Supervised+LLM ensemble models

In all three tasks, LLMs correctly classified cases not correctly classified by specialized models (Supplementary Fig. 5), suggesting that these methods may complement one another. To test this hypothesis, we combined LLM outputs with those of specialized models using ensemble learning (Fig. 2a). We trained logistic regression, XGBoost, and random forest classifiers on prediction results derived from both LLM probability outputs and task-specific model scores (MetaCH, AlphaMissense, GDD-ENS) and assessed the performance of these classifiers using 10-fold cross validation. Ensemble approaches outperformed individual models across all three tasks (p<0.001, Mann-Whitney U test). For distinguishing cancer vs CHIP mutations, the logistic regression ensemble combining Qwen3-30B and MetaCH features achieved F1 0.924 compared to 0.898 for Qwen3-30B alone and 0.892 for MetaCH alone (+3.5%) (Fig. 2a). Similar improvements were observed for driver annotation (logistic regression ensemble combining GPT-5 and AlphaMissense; F1 0.897 vs. 0.866 for AlphaMissense, +3.6%; Fig. 2b) and cancer type prediction (logistic regression ensemble combining GPT-5 and GDD-ENS; F1 0.601 vs. 0.570 for GDD-ENS, +5.3%; Fig. 2c). To test generalizability of these models to out-of-distribution datasets, we also performed leave-one-center-out cross-validation for cancer type prediction (Fig. 2d). Ensemble models generalized robustly across institutions with an average improvement of 20.4% across centers relative to GDD-ENS (Fig. 2d, Supplementary Table 3). Notably, improvements were concentrated at centers where GDD-ENS performed poorly. For example, at the Netherlands Cancer Institute (NKI), where feature availability was incomplete, incorporating LLM outputs increased performance by up to 52.8%. These results suggest that generalist LLM outputs can improve the predictive power of a specialized model in data that is out-of-distribution for the latter.

**Figure 2.**
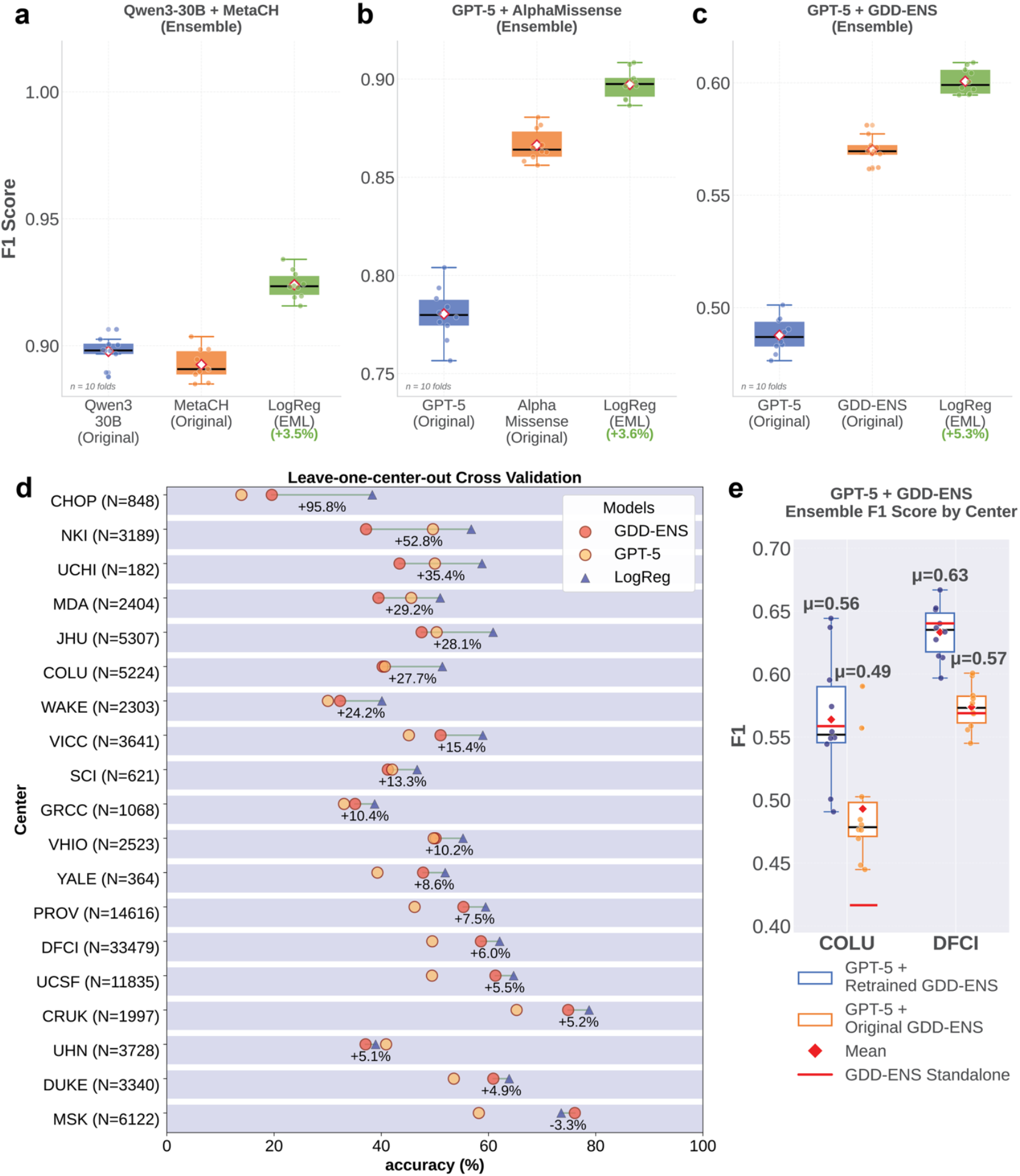
Ensemble model performance cross validation. a-c, F1 performance (10-fold cross-validation) of the logistic regression ensemble (LogReg/EML, green) compared with baseline LLM and specialized models across three tumor genomics tasks: mutation status (tumor-somatic vs. CHIP; Qwen3-30B + MetaCH; a), variant oncogenicity (benign vs. oncogenic; GPT-5 + AlphaMissense; b), and cancer type prediction (GPT-5 + GDD-ENS; c). Percent improvement is shown in green, relative to the better baseline. d, Leave-one-center-out cross-validation for cancer type prediction. Centers are ranked by percent accuracy improvement from GDD-ENS to the best-performing ensemble (logistic regression). e, F1 score (10-fold cross-validation) of GPT-5 + GDD-ENS ensembles for cancer type prediction at Columbia University (COLU) and Dana-Farber Cancer Institute (DFCI), using the original GDD-ENS (orange) versus GDD-ENS retrained on institution-specific data (blue).

To test the impact of GDD-ENS retraining on a more representative dataset for out-of-distribution data, we retrained GDD-ENS on institution-specific data from Dana-Farber Cancer Institute (DFCI) and Columbia University (COLU), the two largest cohorts after MSK (Fig. 2e). Retraining on institution-specific data yielded performance gains at both sites that could not be improved by further ensembling with zero-shot LLM results (Fig. 2e). Thus, for the challenging task of cancer type inference, fine-tuning on institution-specific data currently yields best-in-class performance, when sufficient data for such retraining are available.

## Discussion

LLMs have already disrupted the clinical landscape in traditionally human-oriented tasks including documentation, diagnostic reasoning, and patient communication^20^. Tumor genomics interpretation represents a particularly challenging set of clinically important tasks typically done by bioinformatics algorithms. The value of LLMs in this setting is less widely studied and it is unclear whether their training corpus contains useful information for these tasks. Here, we find that general-purpose LLMs achieve performance equal to specialized models for distinguishing tumor from non-tumor variants. For identifying driver mutations and inferring cancer type, specialized models currently retain an advantage, although the rapid improvement of zero-shot LLMs for these challenging tasks suggests that LLMs may eventually equal or improve upon specialized models.

Ensemble methods integrating LLM outputs with those of task-specific models improved performance across all evaluated tasks, suggesting that the broad pretraining of LLMs provides useful complementary information to help with variant interpretation. In the case of cancer type inference, retraining a supervised model on initially out-of-distribution data abrogated the benefit of an ensemble model, although the practical challenges associated with retraining and even implementing specialized models may make such retraining infeasible for many users.

This study has limitations. Ground-truth labels for variant oncogenicity relied on curated knowledge bases, which are necessarily incomplete and may evolve over time. The analyzed cohorts are enriched for common cancer types and demographics. It was not feasible to assess the full and rapidly expanding LLM landscape. Nonetheless, our study suggests the growing utility of LLMs for complex tumor genomics tasks typically performed by tailor-made models and supports the clinical use of these models alongside current methods.

## Methods

### Datasets and cohort selection

For the tumor-somatic versus CHIP classification task, we assembled a dataset that combined tumor-somatic mutations from MSK patients in the AACR Project GENIE v17.0 public release with CHIP variants matched at the sample level from a previously published MSK cohort^11^. The evaluation set was drawn from both sources, with an approximate 2:1 ratio of tumor-somatic to CHIP variants, and excluded any non-zero VAF variants that overlapped with MetaCH’s training data. The final evaluation cohort comprised 11,908 patients and 15,769 samples, with 34,415 variants labeled as tumor-somatic (n = 22,900) or CHIP (n = 11,515).

For oncogenic classification, we used MSK-IMPACT data with ground-truth labels derived from OncoKB^13^, COSMIC, and dbSNP annotations. The evaluation set was restricted to missense variants and constructed to maintain an approximate 2:1 ratio of oncogenic to benign variants. The final evaluation cohort included 10,489 patients and 10,752 samples, with 13,469 variants labeled as oncogenic (n = 8,817) or benign (n = 4,652).

For cancer type prediction, we used data from the AACR Project GENIE consortium comprising 97,074 patients and 102,791 samples spanning 34 cancer types, consistent with the GDD-ENS training set definitions. Any samples included in GDD-ENS training were removed from the final evaluation set. Genomic features mirrored those available in GENIE and included somatic mutations, somatic copy-number alterations, chromosomal arm-level changes, and gene fusions.

### Large language model evaluation

We evaluated multiple state-of-the-art LLMs including Claude Opus 4.5, Claude Sonnet 4.5, Claude Sonnet 3.7, GPT-5, GPT-4o, o3-mini, DeepSeek-R1, DeepSeek-R1-Distill-Qwen-32B, Qwen3-235B, Qwen3-30B-A3B-Instruct-2507, and MedGemma-27B. All models were evaluated in a zero-shot setting without task-specific fine-tuning. Structured prompts were designed to present genomic information and request predictions in a standardized JSON format, including prediction labels, probability estimates, and explanatory text.

For Task 1 (tumor-somatic vs. CHIP), prompts included gene name, exon, mutation type, amino acid change, variant allele fraction, cancer type, and detailed cancer type. For Task 2 (oncogenicity classification), prompts included gene name, exon, mutation type, amino acid change, and cancer type. For Task 3 (cancer type prediction), prompts included patient sex, somatic mutations, copy number alterations, chromosomal arm-level changes, and fusion events.

### Baseline supervised models

MetaCH ^5^ is a machine learning framework for classifying the origin of cell-free DNA variants, trained on MSK-IMPACT and msk_ch_2020 datasets using features including reference allele, tumor alleles, variant type, gene symbol, gene name, reference genome, VAF, and cancer type.

AlphaMissense^6^ is a deep learning model predicting pathogenicity of missense variants by integrating AlphaFold-derived 3D structure context, protein language modeling of sequence patterns, and training on ClinVar annotations.

GDD-ENS ^7^ is an ensemble deep learning model trained on genomic features for cancer type prediction using MSK-IMPACT and GENIE consortium data.

### Performance evaluation

Model performance was assessed using standard classification metrics including accuracy, F1 score, precision and recall. For cancer type prediction, both top-1 and top-2 accuracy were evaluated. Confusion matrices were generated to visualize classification patterns (Supplementary Fig. 6). Calibration curves were constructed to assess probability and estimate reliability (Supplementary Fig. 7).

### Ensemble model development

For cancer type prediction, we developed an ensemble approach using logistic regression, XGBoost and random forest combining predictions from LLM and each task-specific model (MetaCH, AlphaMissense and GDD-ENS) with corresponding probability scores. The ensemble incorporated the top two predictions and associated probabilities from both LLM and task-specific models to generate a final cancer type prediction. Ensemble performance was compared against individual model performance using the same evaluation metrics (Supplementary Table 4). Leave-one-center-out cross-validation was performed to assess generalizability across GENIE consortium sites (Supplementary Table 3).

### GDD-ENS retraining

GDD-ENS upsampling strategies, feature derivation and selection methods were used to develop center-specific models for DFCI and COLU, with limited adaptations. A minimum of 10 samples per type was required for GDD-ENS architecture compatibility, thus the COLU center model was only trained across 19 cancer types; DFCI had sufficient samples to retrain across the original 38 types. Additionally, DFCI did not report arm-level copy number features, and COLU did not report any gene or arm-level copy number, or fusions. Unlike GDD-ENS, there were no genomic content thresholds used to subset sample quality during training or testing.

Finally, as not all DFCI or COLU genomic panels perform tumor-normal matched sequencing, all detected mutations were reported and incorporated into center specific feature tables regardless of somatic status. Center-specific models using GDD-ENS architecture and hyperparameter optimization techniques were then trained and tested, and results were reported analogously to prior methods.

### Interpretability analysis

LLM feature attributions were extracted from the structured output format, which included key genes, chromosomal arm changes, somatic copy number alterations, and gene fusions identified as influential for predictions. SHAP (SHapley Additive exPlanations) ^21^ values were computed for the GDD-ENS model to identify feature importances. Correlation between LLM-attributed features and SHAP importances was assessed using Pearson correlation.

### Survival analysis

Kaplan-Meier survival analysis was performed to assess the clinical relevance of LLM oncogenicity predictions for variants of uncertain significance (VUS). 12,252 patients with NSCLC were stratified by KEAP1 mutation status and LLM-predicted oncogenicity^4,14,22^. Log-rank tests were used to compare survival curves between groups.

### Statistical analysis

All statistical analyses were performed using Python 3.10 with NumPy, Pandas, Scikit-learn libraries and R 4.4 with survival and survminer packages. Mann-Whitney U tests were computed using bootstrap resampling with 1,000 iterations. Statistical significance was assessed at α = 0.05.

## Data Availability

The MSK-GENIE dataset is publicly available through the AACR Project GENIE data portal (https://www.aacr.org/professionals/research/aacr-project-genie/). The msk_ch_2023 dataset is available from the cBioPortal (https://www.cbioportal.org/study/summary?id=msk_ch_2023).

OncoKB annotations are available at https://www.oncokb.org/. COSMIC annotations (CancerMutationCensus_AllData_v102_GRCh37.tsv) are available at https://cancer.sanger.ac.uk/cmc/. dbSNP annotations (common_no_known_medical_impact_20170905.vcf.gz) are available at https://ftp.ncbi.nlm.nih.gov/pub/clinvar/vcf_GRCh37/archive_1.0/2017/.

## Code Availability

Code for LLM prompting, evaluation pipelines, and statistical analyses is available on GitHub: https://github.com/smilejennyyu/llm_cancer_type_classification/tree/main.

## Acknowledgements

We acknowledge the contributions of the AACR Project GENIE consortium and all participating institutions. This work was supported by K08CA286842 and P30 CA008748.

## Author Contributions

J.Y. and J.J. designed and performed experiments, analyzed data, and wrote the manuscript. M.D., M.W., T.N.T., C.F., L.M. and K.U. contributed to data processing and analysis. M.W. and J.C. contributed to setting up LLMs. M.B. and N.S. provided access to MSK-IMPACT data and contributed to study design. R.L. provided expertise on clonal hematopoiesis. Q.M. and J.J. conceived the study, conceptualized the project and supervised the research. All authors edited the manuscript, reviewed and approved the paper.

