## supplementary figures for "Generalist large language models complement tailor-made predictors for tumor genomics interpretation"

#### Supplementary Information Table of Contents

**1. Figures: Figures S1–7**

**2. Tables: Tables S1–4**

### Supplementary Figures and Tables

#### 1. Figures

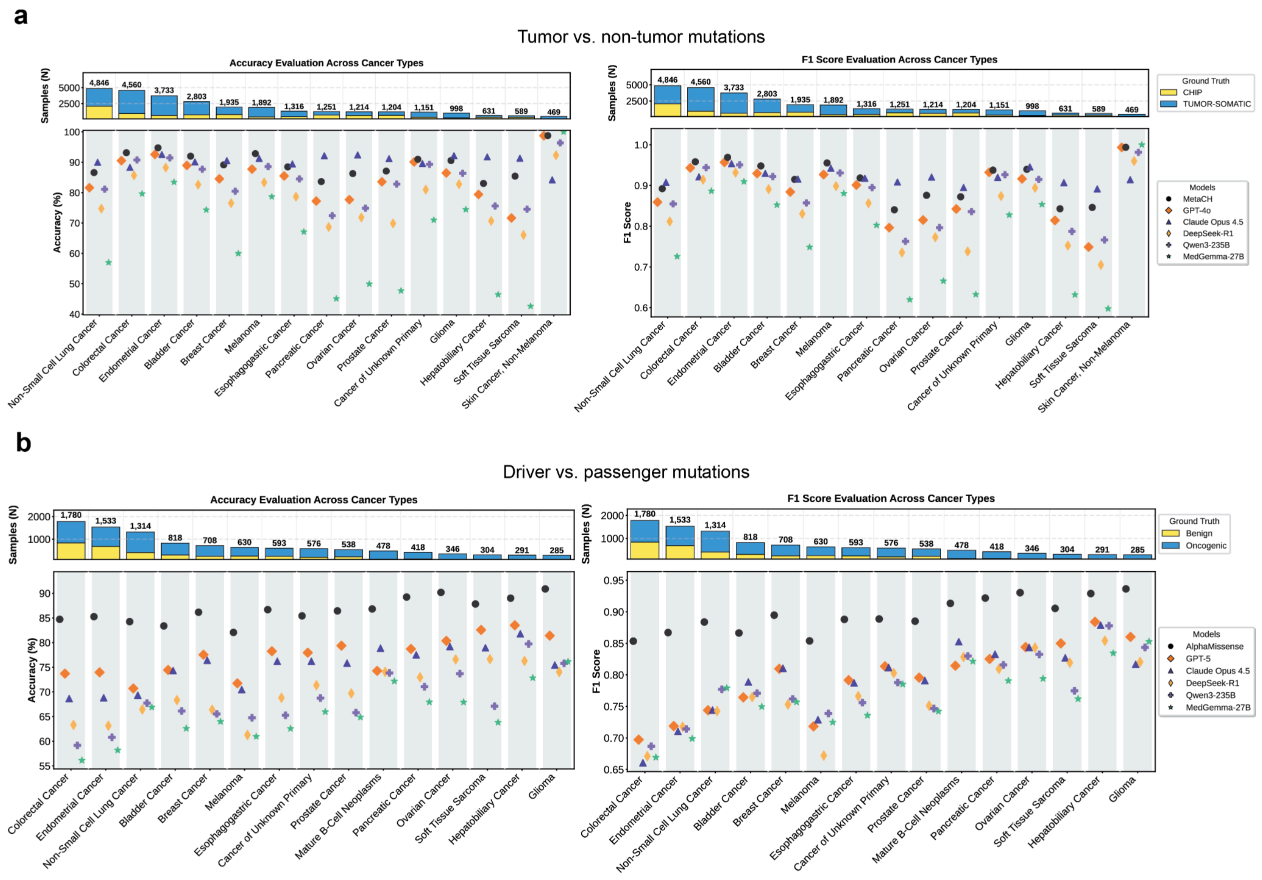


#### Figure S1 | Performance stratified by cancer type

**a)** Tumor vs. non-tumor mutation classification stratified by cancer type. Top, sample counts and class composition (CHIP vs. tumor-somatic). Bottom, per-cancer accuracy (left) and F1 score (right) for MetaCH and LLMs. **b)** Driver vs. passenger mutation classification stratified by top 15 cancer types. Top, sample counts and class composition (benign vs. oncogenic). Bottom, per-cancer accuracy (left) and F1 score (right) for AlphaMissense and LLMs. Cancer types are ordered by sample size.

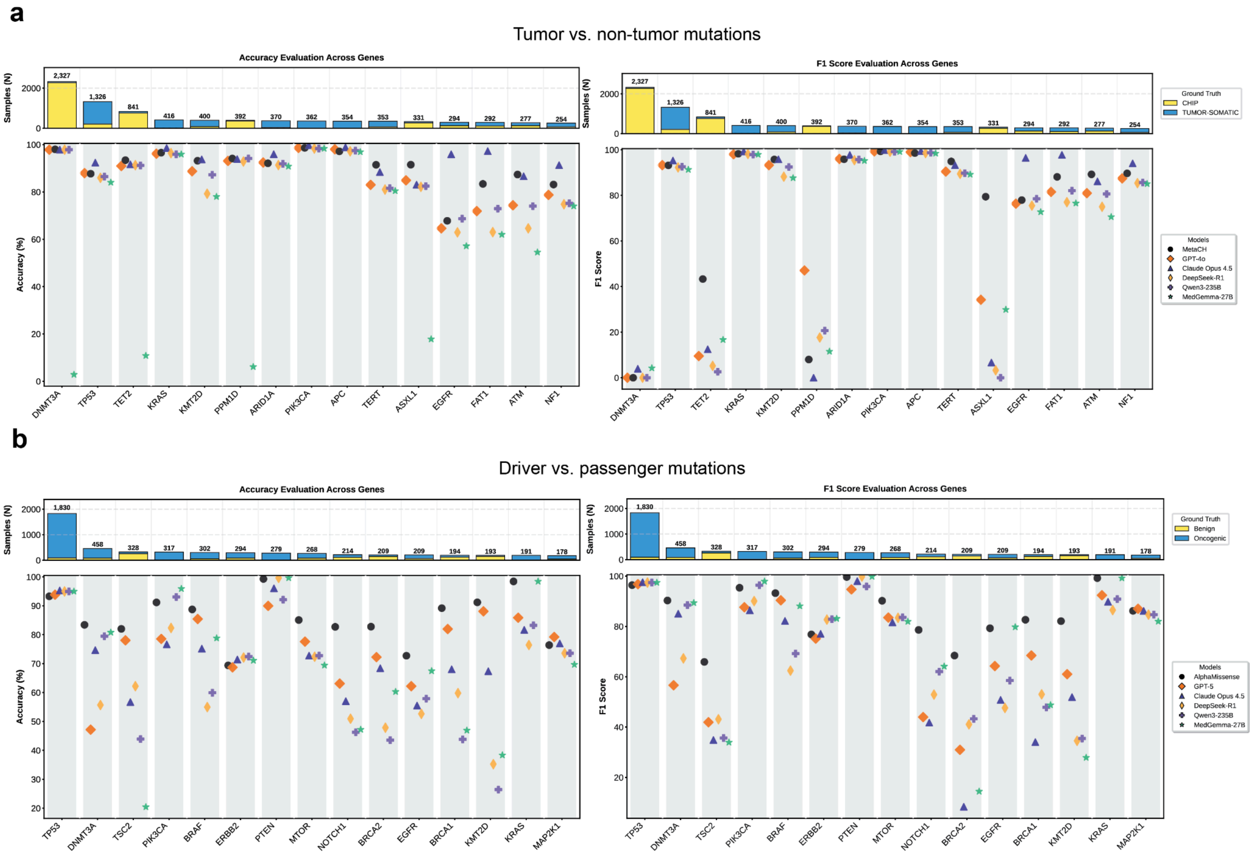


#### Figure S2 | Performance stratified by gene

**a**) Tumor vs. non-tumor mutation classification stratified by gene. Top, sample counts and class composition (CHIP vs. tumor-somatic). Bottom, per-gene accuracy (left) and F1 score (right) for MetaCH and LLMs. **b)** Driver vs. passenger mutation classification stratified by gene. Top, sample counts and class composition (benign vs. oncogenic). Bottom, per-gene accuracy (left) and F1 score (right) for AlphaMissense and LLMs. Genes are ordered by sample size.

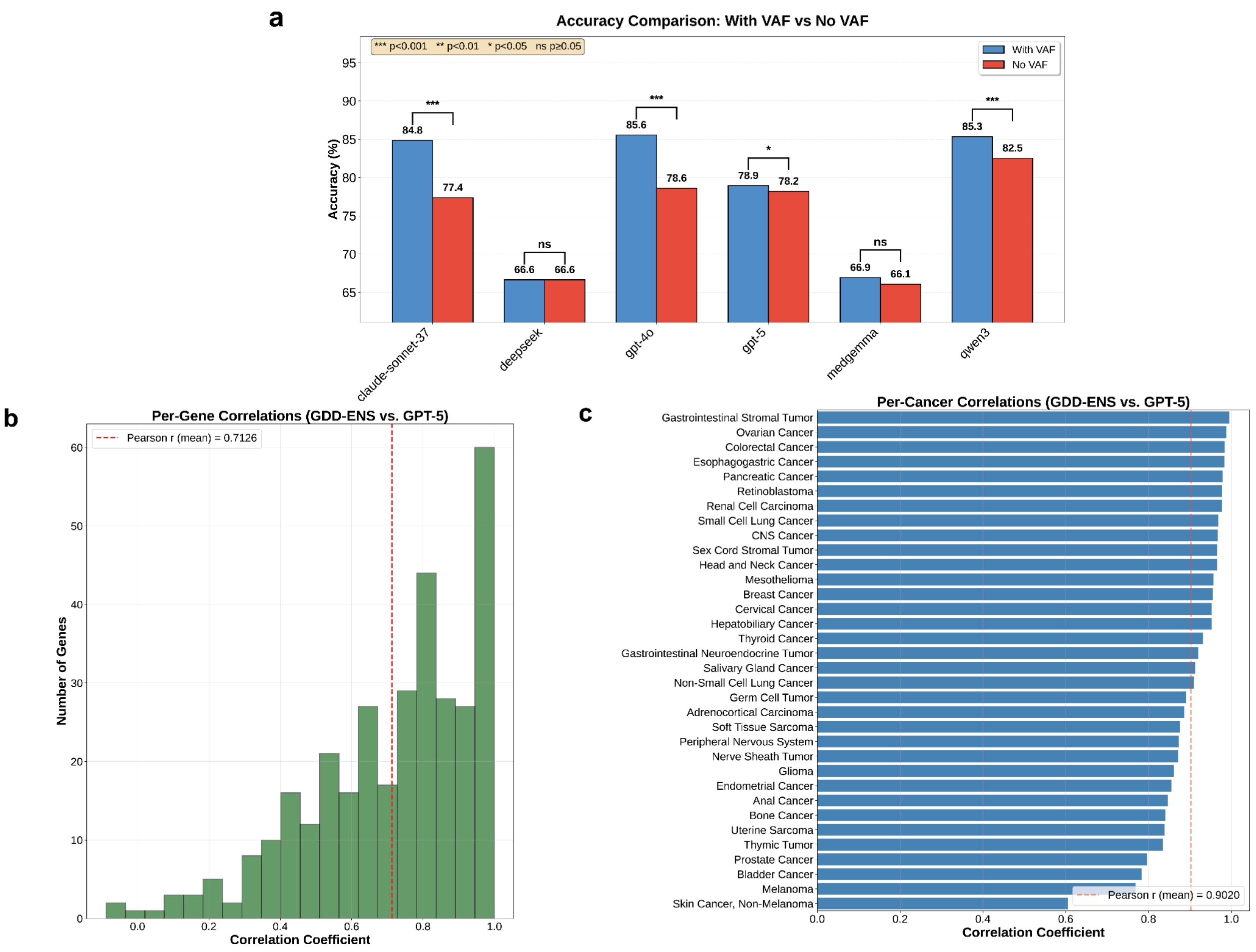


### Figure S3 | Interpretability and ablation analyses

**a)** Accuracy with versus without variant allele fraction (VAF) in prompts for tumor vs. non-tumor mutation classification across LLMs. Bars show mean accuracy; brackets indicate pairwise comparisons (ns, $P$<0.05, *$P$<0.01, **$P$<0.001). **b)** Distribution of per-gene Pearson correlations between GDD-ENS and GPT-5 predictions; dashed line indicates the mean correlation. **c)** Per-cancer Pearson correlations between GDD-ENS and GPT-5 predictions; dashed line indicates the mean correlation.

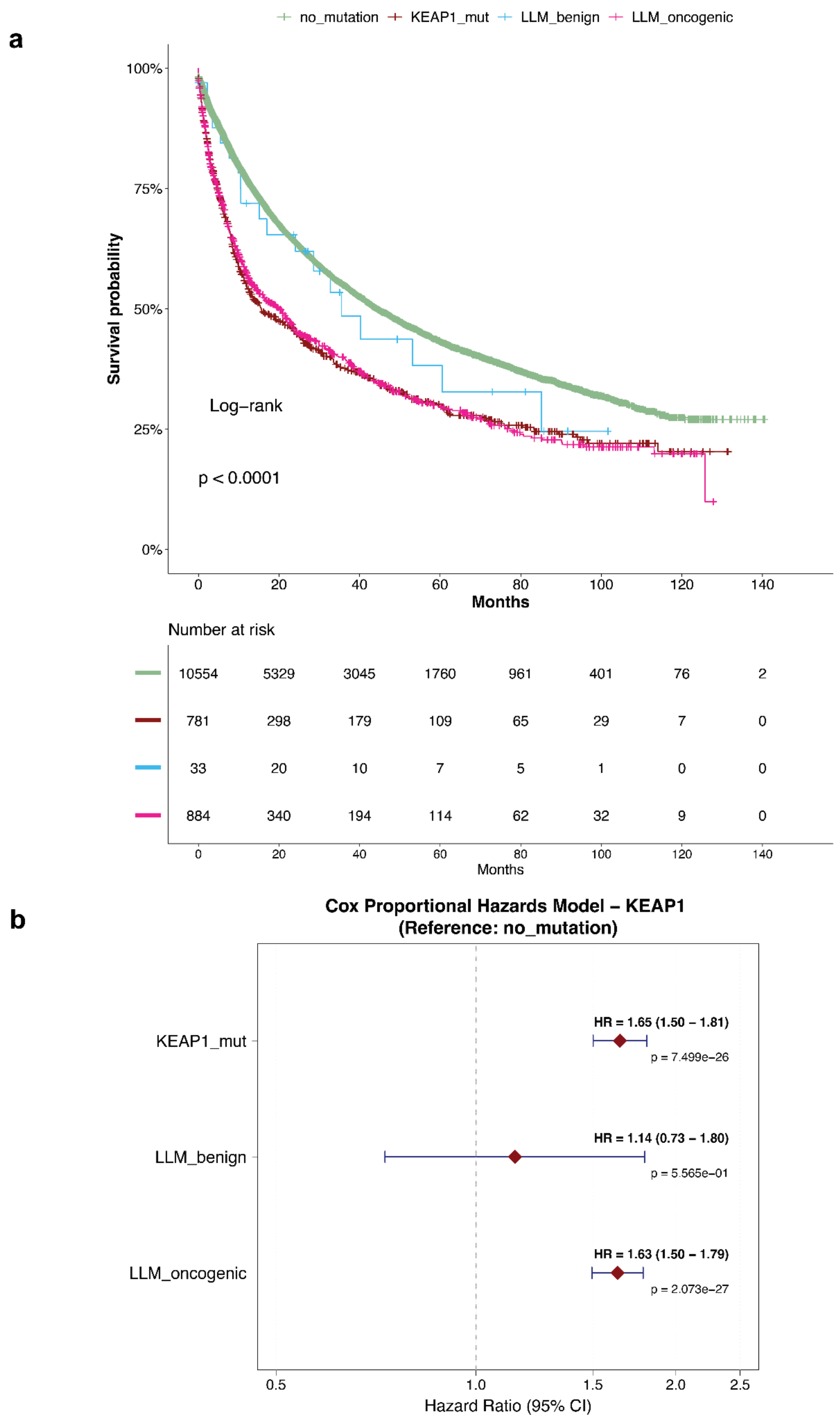


#### Figure S4 | Survival analysis of KEAP1 and LLM-derived labels

**a)** Kaplan–Meier overall survival curves for no mutation, KEAP1-mutant, LLM-benign, and LLM-oncogenic groups, with number-at-risk table and log-rank *P* value. **b)** Cox proportional hazards model (reference, no mutation) showing hazard ratios and 95% confidence intervals for KEAP1-mutant, LLM-benign, and LLM-oncogenic groups.

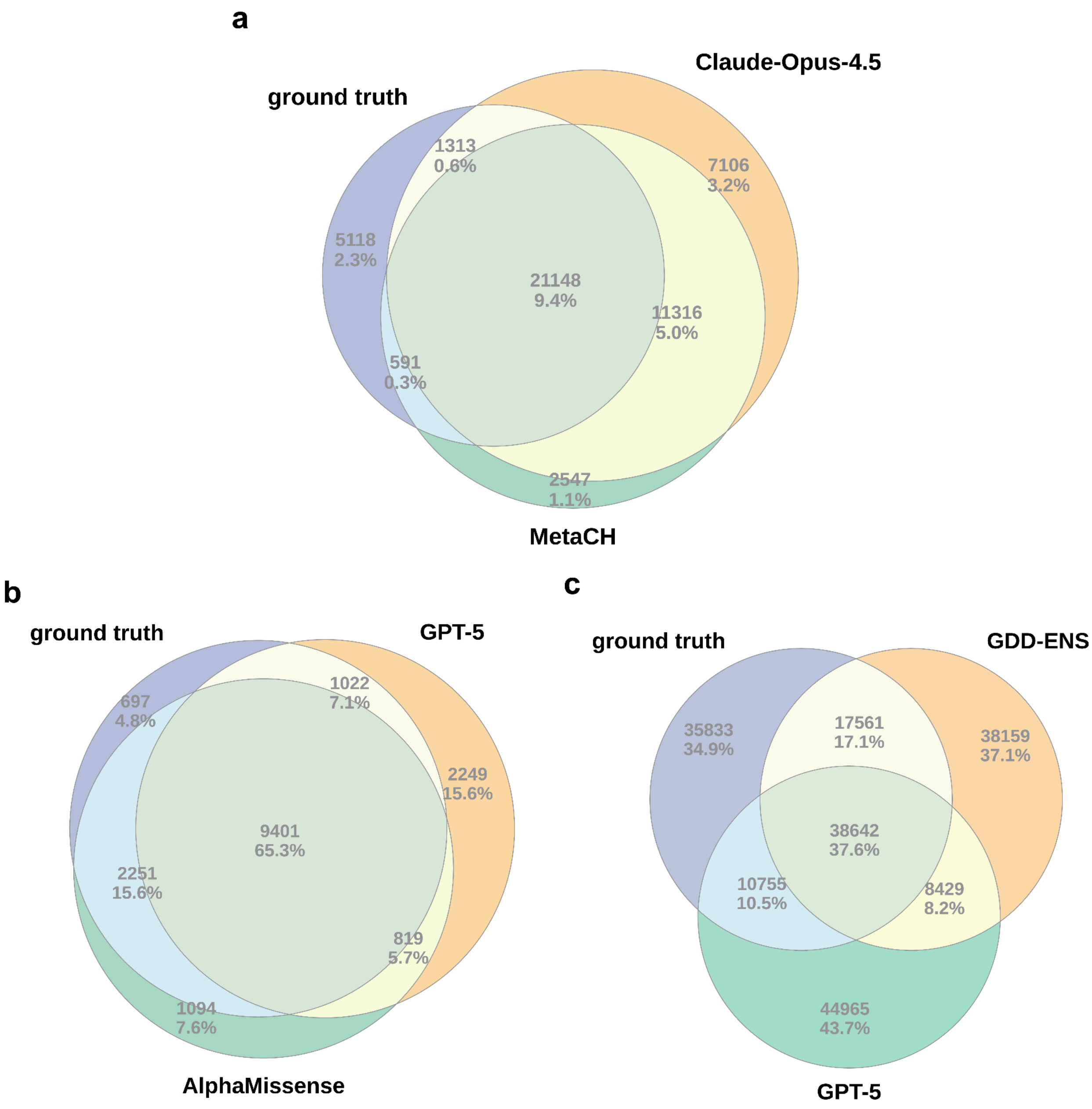


#### Figure S5 | Overlap of correctly classified samples

**a)** Venn diagram of overlap among ground truth, Claude Opus 4.5, and MetaCH for tumor vs. non-tumor mutation classification. **b)** Venn diagram of overlap among ground truth, GPT-5, and AlphaMissense for driver vs. passenger mutation classification. **c)** Venn diagram of overlap among ground truth, GDD-ENS, and GPT-5 for cancer type inference. Numbers indicate counts and percentages.

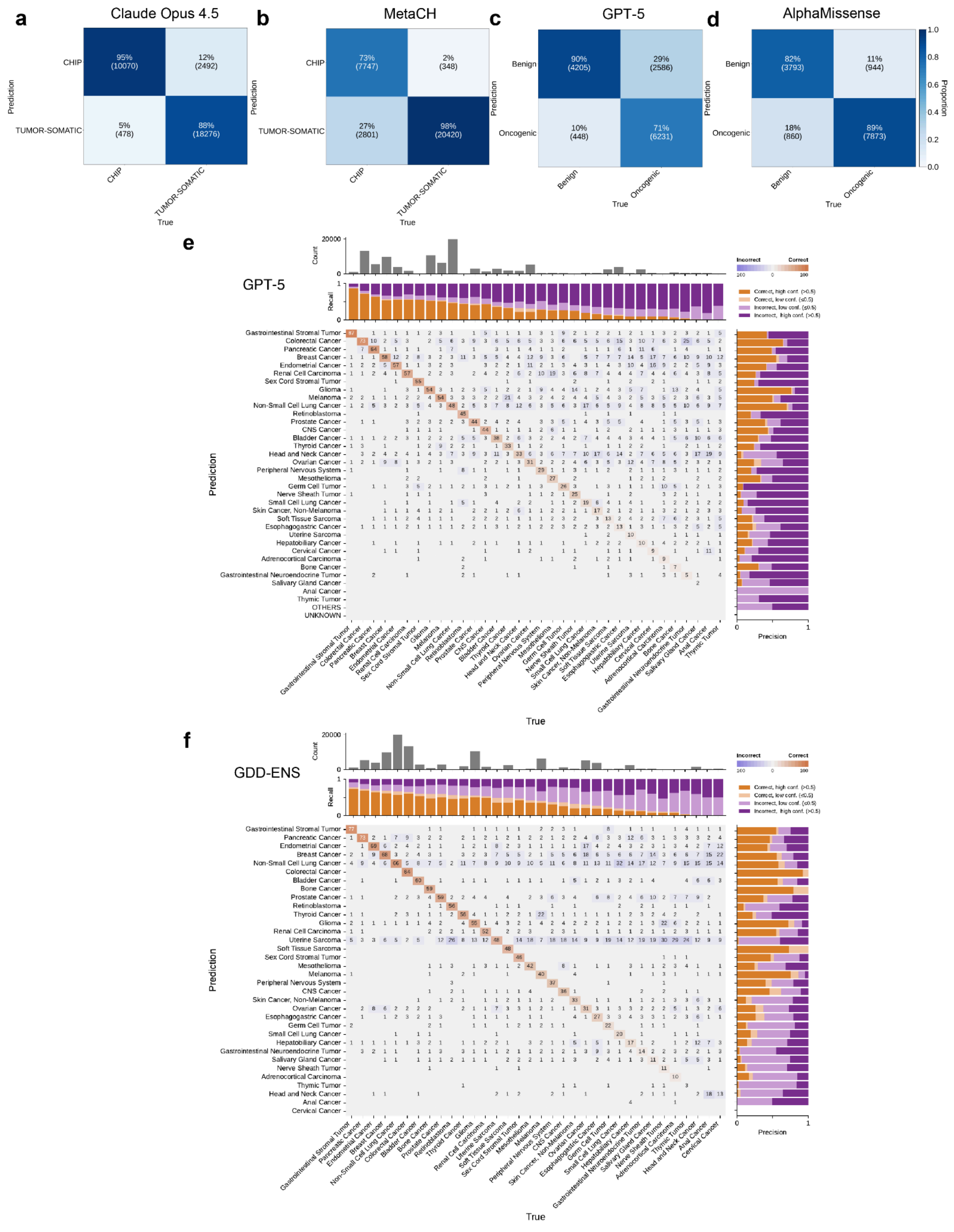


### Figure S6 | Confusion matrices across tasks

**a–b)** Normalized confusion matrices for Claude Opus 4.5 (**a**) and MetaCH (**b**) on tumor vs. non-tumor mutation classification. **c–d)** Normalized confusion matrices for GPT-5 (**c**) and AlphaMissense (**d**) on driver vs. passenger mutation classification. **e–f)** Cancer type inference confusion matrices for GPT-5 (**e**) and GDD-ENS (**f**), with class-wise recall (top) and precision (right). Colors indicate correct vs. incorrect predictions, stratified by confidence threshold categories shown in the legend.

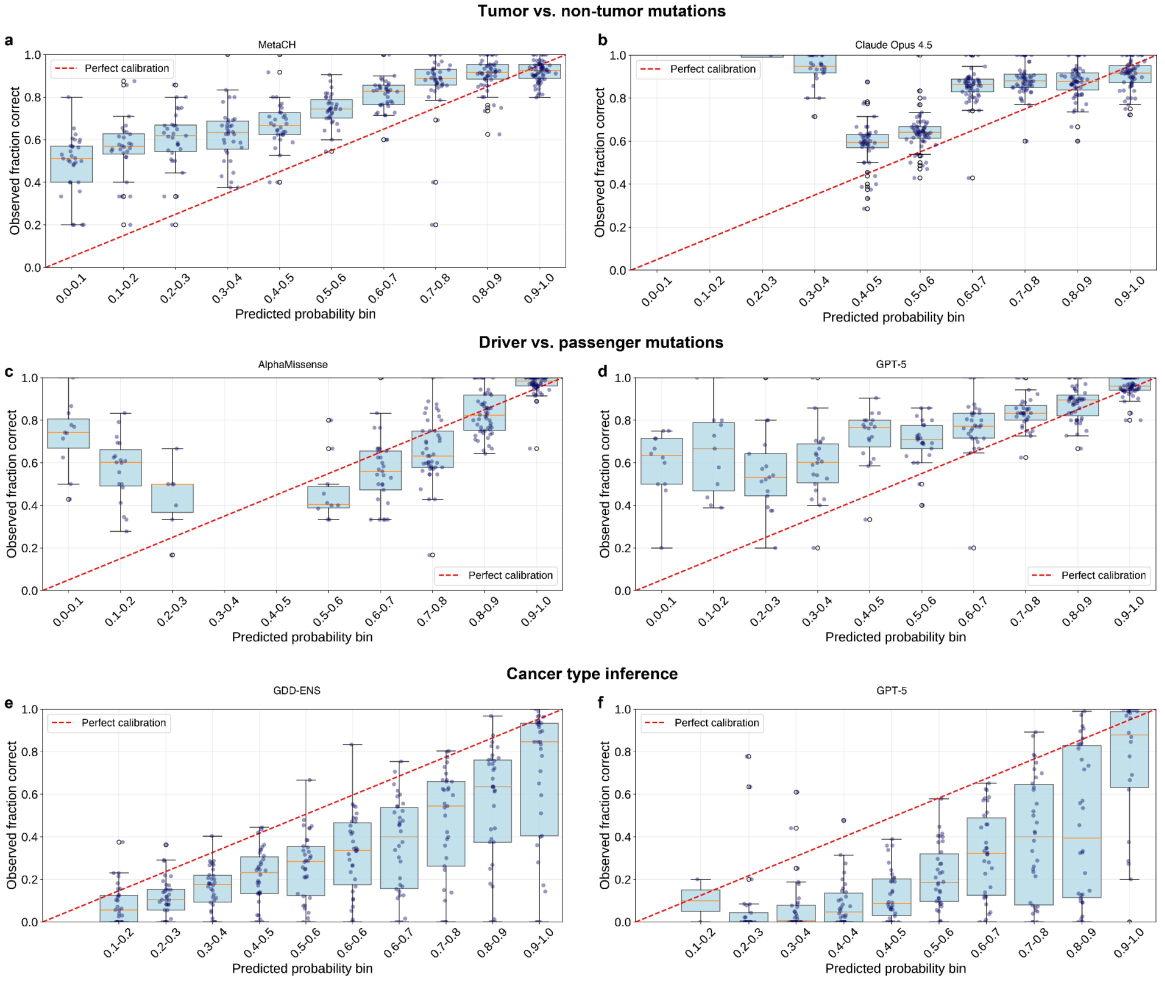


#### Figure S7 | Calibration across tumor-genomic prediction tasks

Calibration plots showing observed fraction correct versus predicted probability bins (dashed line, perfect calibration). **a–b)** MetaCH **(a)** and Claude Opus 4.5 **(b)** for tumor vs. non-tumor mutation classification. **c–d)** AlphaMissense **(c)** and GPT-5 **(d)** for driver vs. passenger mutation classification. **e–f)** GDD-ENS **(e)** and GPT-5 **(f)** for cancer type inference.

#### 2. Tables

**Table S1:** Performance Across Different Models for All Three Tasks

| Task | Model | Accuracy | Top-2 Accuracy | Precision | Recall | F1 |
|---|---|---|---|---|---|---|
| Mutation Status (Tumor-somatic vs. CHIP) | MetaCH | 0.8986 | n/a | 0.8793 | 0.9819 | 0.9278 |
| | o3-mini | 0.7923 | n/a | 0.768 | 0.9864 | 0.8636 |
| | gpt-4o | 0.856 | n/a | 0.8298 | 0.9862 | 0.9013 |
| | gpt-5 | 0.7906 | n/a | 0.7721 | 0.9731 | 0.861 |
| | DeepSeek-R1-Distill-Qwen-32B | 0.6664 | n/a | 0.6664 | 1 | 0.7998 |
| | DeepSeek-R1 | 0.7936 | n/a | 0.7815 | 0.9581 | 0.8608 |
| | Qwen3-30B | 0.8534 | n/a | 0.8245 | 0.9909 | 0.9001 |
| | Qwen3-VL-235B-A22B | 0.8429 | n/a | 0.8202 | 0.9789 | 0.8925 |
| | medgemma-27b-text-it | 0.6693 | n/a | 0.6684 | 0.9999 | 0.8012 |
| | claude-sonnet-3.7 | 0.8486 | n/a | 0.8388 | 0.9565 | 0.8938 |
| | claude-sonnet-4.5 | 0.8587 | n/a | 0.9472 | 0.8345 | 0.8873 |
| | **claude-opus-4.5** | **0.9038** | n/a | **0.9747** | **0.8785** | **0.9241** |
| Oncogenicity (Benign vs. Oncogenic) | AlphaMissense | 0.8661 | n/a | 0.9015 | 0.8929 | 0.8972 |
| | o3-mini | 0.7521 | n/a | 0.7868 | 0.8522 | 0.8182 |
| | gpt-4o | 0.6939 | n/a | 0.8041 | 0.7037 | 0.7506 |
| | **gpt-5** | **0.7747** | n/a | **0.9329** | **0.7067** | **0.8042** |
| | DeepSeek-R1-Distill-Qwen-32B | 0.6584 | n/a | 0.6584 | 0.9936 | 0.792 |
| | DeepSeek-R1 | 0.6991 | n/a | 0.7503 | 0.81 | 0.779 |
| | Qwen3-30B | 0.6538 | n/a | 0.6558 | 0.9915 | 0.7894 |
| | Qwen3-VL-235B-A22B | 0.683 | n/a | 0.708 | 0.8777 | 0.7838 |
| | medgemma-27b-text-it | 0.659 | n/a | 0.6862 | 0.8828 | 0.7722 |
| | claude-sonnet-3.7 | 0.7311 | n/a | 0.8012 | 0.7837 | 0.7924 |
| | claude-sonnet-4.5 | 0.6773 | n/a | 0.7774 | 0.7105 | 0.7424 |
| | claude-opus-4.5 | 0.7514 | n/a | 0.8523 | 0.7502 | 0.798 |
| Cancer Type (34 types) | GDD-ENS | 0.5468 | 0.6585 | 0.6361 | 0.5468 | 0.5702 |
| | o3-mini | 0.4342 | 0.5518 | 0.4716 | 0.4342 | 0.4218 |
| | gpt-4o | 0.3468 | 0.4708 | 0.3792 | 0.3468 | 0.3209 |
| | **gpt-5** | **0.4806** | **0.5973** | **0.5229** | **0.4806** | **0.4876** |
| | DeepSeek-R1-Distill-Qwen-32B | 0.2401 | 0.3428 | 0.3627 | 0.2401 | 0.2289 |
| | DeepSeek-R1 | 0.4051 | 0.5218 | 0.4567 | 0.4051 | 0.4035 |
| | Qwen3-30B | 0.2534 | 0.3477 | 0.3473 | 0.2534 | 0.2564 |
| | Qwen3-VL-235B-A22B | 0.3344 | 0.435 | 0.378 | 0.3344 | 0.3106 |
| | medgemma-27b-text-it | 0.209 | 0.281 | 0.3219 | 0.209 | 0.1933 |
| | claude-sonnet-3.7 | 0.4418 | 0.5634 | 0.4528 | 0.4418 | 0.4133 |
| | claude-sonnet-4.5 | 0.4311 | 0.5268 | 0.4575 | 0.4311 | 0.4098 |
| | claude-opus-4.5 | 0.4803 | 0.5972 | 0.4933 | 0.4803 | 0.465 |

**Table S2:** Per-cancer Performance Across Different Models for All Three Tasks (F1)

| Task | Model | Q1 | Median | Q3 | IQR |
|---|---|---|---|---|---|
| Mutation Status (Tumor-somatic vs. CHIP) | MetaCH | 85.715 | 89.614 | 92.286 | 6.571 |
| | o3-mini | 67.83 | 75.149 | 81.837 | 14.007 |
| | gpt-4o | 77.335 | 81.804 | 88.111 | 10.776 |
| | gpt-5 | 68.001 | 75.097 | 81.641 | 13.64 |
| | DeepSeek-R1-Distill-Qwen-32B | 47.498 | 59.819 | 73.783 | 26.285 |
| | DeepSeek-R1 | 69.543 | 74.773 | 81.222 | 11.678 |
| | Qwen3-30B | 78.465 | 82.009 | 87.818 | 9.354 |
| | Qwen3-VL-235B-A22B | 74.915 | 80.675 | 85.983 | 11.068 |
| | medgemma-27b-text-it | 48.101 | 59.974 | 74.415 | 26.313 |
| | claude-sonnet-3.7 | 74.625 | 81.535 | 86.367 | 11.741 |
| | claude-sonnet-4.5 | 83.649 | 87.946 | 90.185 | 6.536 |
| | **claude-opus-4.5** | **88.361** | **91.89** | **93.997** | **5.635** |
| Oncogenicity (Benign vs. Oncogenic) | AlphaMissense | 86.047 | 89.582 | 92.69 | 6.643 |
| | o3-mini | 75.104 | 81.347 | 88.076 | 12.972 |
| | gpt-4o | 70.641 | 73.937 | 83.553 | 12.911 |
| | gpt-5 | **79.203** | **83.827** | **90.135** | **10.932** |
| | DeepSeek-R1-Distill-Qwen-32B | 68.053 | 74.884 | 83.028 | 14.976 |
| | DeepSeek-R1 | 71.163 | 78.306 | 83.999 | 12.836 |
| | Qwen3-30B | 65.614 | 74.398 | 82.095 | 16.481 |
| | Qwen3-VL-235B-A22B | 68.496 | 78.155 | 83.999 | 15.504 |
| | medgemma-27b-text-it | 66.493 | 74.931 | 83.121 | 16.628 |
| | claude-sonnet-3.7 | 73.037 | 79.512 | 88.036 | 14.999 |
| | claude-sonnet-4.5 | 68.9 | 75.996 | 84.408 | 15.507 |
| | claude-opus-4.5 | 76.221 | 82.299 | 88.807 | 12.586 |
| Cancer Type (34 types) | GDD-ENS | 17.861 | 40.922 | 58.529 | 40.669 |
| | o3-mini | 3.93 | 14.477 | 45.716 | 41.785 |
| | gpt-4o | 1.184 | 7.004 | 29.011 | 27.827 |
| | gpt-5 | **10.738** | **29.998** | **52.327** | **41.589** |
| | DeepSeek-R1-Distill-Qwen-32B | 0.119 | 3.974 | 15.264 | 15.145 |
| | DeepSeek-R1 | 3.976 | 14 | 41.008 | 37.031 |
| | Qwen3-30B | 0.535 | 4.47 | 29.772 | 29.237 |
| | Qwen3-VL-235B-A22B | 0.702 | 4.264 | 25.785 | 25.083 |
| | medgemma-27b-text-it | 0.007 | 0.598 | 13.53 | 13.523 |
| | claude-sonnet-3.7 | 2.176 | 9.73 | 49.565 | 47.389 |
| | claude-sonnet-4.5 | 1.185 | 10.524 | 39.853 | 38.668 |
| | claude-opus-4.5 | 3.712 | 14.631 | 46.437 | 42.725 |

**Table S3:** Ensemble Models Performance Summary (Leave-One-Center-Out Cross Validation)

| Method | Accuracy (mean±sd) | F1 (mean±sd) | Precision (mean±sd) | Recall (mean±sd) |
|---|---|---|---|---|
| GDD-ENS Original | 0.479±0.143 | 0.518±0.143 | **0.666±0.136** | 0.479±0.143 |
| LogReg | 0.553±0.114 | 0.567±0.123 | 0.630±0.138 | 0.553±0.114 |
| Random Forest | 0.552±0.123 | 0.562±0.128 | 0.630±0.141 | 0.552±0.123 |
| XGBoost | **0.561±0.124** | **0.577±0.126** | 0.647±0.133 | **0.561±0.124** |
| GPT-5 Original | 0.449±0.111 | 0.480±0.123 | 0.587±0.140 | 0.449±0.111 |

**Table S4:** Ensemble Models Performance Summary (10-fold Cross Validation)

| Task | Method | Accuracy (mean±sd) | F1 (mean±sd) | Precision (mean±sd) | Recall (mean±sd) |
|---|---|---|---|---|---|
| Mutation status | MetaCH | 0.899±0.005 | 0.895±0.005 | 0.904±0.004 | 0.899±0.005 |
| | claude-opus-4.5 | 0.904±0.009 | 0.906±0.009 | 0.916±0.006 | 0.904±0.009 |
| | LogReg | 0.931±0.005 | 0.931±0.005 | 0.931±0.005 | 0.931±0.005 |
| | Random Forest | 0.914±0.008 | 0.914±0.008 | 0.914±0.007 | 0.914±0.008 |
| | **XGBoost** | **0.940±0.005** | **0.941±0.005** | **0.941±0.004** | **0.940±0.005** |
| Oncogenicity | AlphaMissense | 0.866±0.009 | 0.866±0.008 | 0.867±0.008 | 0.866±0.009 |
| | GPT-5 Original | 0.775±0.014 | 0.780±0.013 | 0.825±0.010 | 0.775±0.014 |
| | LogReg | 0.897±0.007 | 0.897±0.007 | 0.898±0.007 | 0.897±0.007 |
| | **Random Forest** | **0.905±0.009** | **0.904±0.009** | **0.905±0.008** | **0.905±0.009** |
| | XGBoost | 0.883±0.008 | 0.882±0.007 | 0.882±0.007 | 0.883±0.008 |
| Cancer type | GDD-ENS Original | 0.547±0.006 | 0.570±0.006 | 0.637±0.005 | 0.547±0.006 |
| | GPT-5 Original | 0.481±0.007 | 0.488±0.008 | 0.523±0.009 | 0.481±0.007 |
| | LogReg | 0.615±0.006 | 0.601±0.006 | 0.601±0.006 | 0.615±0.006 |
| | Random Forest | 0.619±0.005 | 0.599±0.005 | 0.604±0.007 | 0.619±0.005 |
| | **XGBoost** | **0.626±0.006** | **0.614±0.006** | **0.620±0.007** | **0.626±0.006** |
